# The RNA helicase DDX3 induces neural crest by promoting AKT activity

**DOI:** 10.1101/785428

**Authors:** Mark Perfetto, Xiaolu Xu, Natasha Yousaf, Jiejing Li, Shuo Wei

**Author notes:** Correspondence should be addressed to: Shuo Wei, Department of Biological Sciences, University of Delaware.

## Abstract

Mutations in the RNA helicase DDX3 have emerged as a frequent cause of intellectual disability in humans. Because many patients carrying DDX3 mutations have additional defects in craniofacial structures and other tissues containing neural crest (NC)-derived cells, we hypothesized that DDX3 is also important for NC development. Using *Xenopus tropicalis* as a model, we show that DDX3 is required for normal NC induction and craniofacial morphogenesis by regulating AKT kinase activity. Depletion of DDX3 decreases AKT activity and AKT-dependent inhibitory phosphorylation of GSK3β, leading to reduced levels of β-catenin and Snai1, two GSK3β substrates that are critical for NC induction. DDX3 function in regulating these downstream signaling events during NC induction is likely mediated by RAC1, a small GTPase whose translation depends on the RNA helicase activity of DDX3. These results suggest an evolutionarily conserved role of DDX3 in NC development by promoting AKT activity, and provide a potential mechanism for the NC-related birth defects displayed by patients harboring mutations in DDX3 and its downstream effectors in this signaling cascade.

## Introduction

Neural crest (NC) cells are multipotent stem cells that are induced to form at the neural plate border (the border of neural plate and epidermis; NPB) during vertebrate gastrulation. After the induction and subsequent maintenance, NC cells migrate out of the closing neural tube to specific destinations, where they differentiate into various types of cells and contribute to many tissues, such as the craniofacial structures, dental tissues, the peripheral nervous system, pigment cells, and cardiac tissues (Bronner and Simões-Costa, 2016; Simões-Costa and Bronner, 2015; Stuhlmiller and Garcia-Castro, 2012). In humans, impaired NC development can lead to birth defects that are collectively called neurocristopathies, including craniofacial disorders, congenital heart diseases, and pigment defects (Dubey and Saint-Jeannet, 2017; Vega-Lopez et al., 2018).

Nearly all of our knowledge on the induction of NC was obtained from studies using *Xenopus* and other non-mammalian vertebrates such as chicks, due to technical difficulties that have been encountered in mice (Barriga et al., 2015; Betters et al., 2018). These studies have uncovered a two-stage process for NC induction. First, the interplay of multiple signaling pathways activates the expression of transcription factors such as Pax3, Zic1 and Msx1 (the “NPB specifiers”), which induce NPB formation. These NPB specifiers subsequently induce other transcription factors, including Snai2, FoxD3 and Sox9 (the “NC specifiers”), to specify the NC fate within the NPB (Bronner and Simões-Costa, 2016; Simões-Costa and Bronner, 2015; Stuhlmiller and Garcia-Castro, 2012). A major signaling pathway that induces the NC in non-mammalian vertebrates is canonical Wnt/β-catenin signaling (hereinafter referred to as “Wnt signaling”), which is required for both the initial formation of the NPB and the subsequent specification of the NC (Li et al., 2018; Prasad et al., 2019). Similarly, the induction of NC from human embryonic stem (ES) cells or induced pluripotent stem (iPS) cells also requires the activation of Wnt signaling (Gomez et al., 2019; Leung et al., 2016; Menendez et al., 2011; Mica et al., 2013), indicating that Wnt function in NC induction is conserved during vertebrate evolution.

The DEAD/H box RNA helicase DDX3 plays important roles in many cellular processes including cell cycle regulation, differentiation, survival, and apoptosis, primarily by regulating ATP-dependent RNA metabolism, such as RNA splicing, nuclear export, and translation (He et al., 2018; Heerma van Voss et al., 2017; Sharma and Jankowsky, 2014). DDX3 contains a helicase ATP-binding domain and a helicase C-terminal domain, both of which are evolutionarily conserved and required for its RNA helicase activity. There are two functional *DDX3* homologs in humans, *DDX3X* and *DDX3Y*, which are localized on the X and Y chromosome, respectively. Although the two gene products share high sequence similarity, they have distinct expression patterns and function. DDX3Y is translated specifically in the testes and is necessary for male fertility, whereas DDX3X is expressed ubiquitously and accounts for most, if not all, DDX3 function in somatic tissues (Kotov et al., 2017; Snijders Blok et al., 2015). In rodents, there is at least one additional homolog of DDX3 that is actively expressed. The tissue specificity of DDX3X and DDX3Y is also different in rodents and primates, suggesting that these genes may have evolved differently in the two mammalian orders (Chang and Liu, 2010; Matsumura et al., 2019). In contrast, *Xenopus tropicalis* has only one *ddx3* homolog, which is localized on an autosome, and the encoded protein is 91% identical to human DDX3X. Similar to human *DDX3X*, the *X. tropicalis ddx3* mRNA is expressed ubiquitously throughout early development (Cruciat et al., 2013). Thus, *X. tropicalis* may be a suitable model to study DDX3 function in somatic tissue development.

Recently, *de novo* mutations in *DDX3X* were reported to be a common cause of intellectual disability in humans (https://ddx3x.org)(Dikow et al., 2017; Kellaris et al., 2018; Lennox et al.; Nicola et al., 2019; Snijders Blok et al., 2015; Wang et al., 2018). These mutations were found primarily in females, and there are strong indications that some of them lead to embryonic lethality in males (Nicola et al., 2019; Snijders Blok et al., 2015). Most of these mutations were predicted to result in decreased DDX3X activity, and the defects in female patients may be caused by haploinsufficiency or dominant-negative effects of the mutants; partial X inactivation of the *DDX3X* gene may also contribute to the phenotypes (Garieri et al., 2018; Lennox et al.; Snijders Blok et al., 2015). Patients with *DDX3X* mutations display various defects in central nervous system (CNS) development, including intellectual disability, autism spectrum disorder, microcephaly, and corpus callosum hypoplasia. Most patients also have craniofacial disorders, ranging from mild dysmorphic facial features to severe oral-facial clefts (Dikow et al., 2017; Kellaris et al., 2018; Lennox et al.; Nicola et al., 2019; Snijders Blok et al., 2015; Wang et al., 2018). Other common symptoms include congenital heart diseases, pigment defects, and hearing/visual impairment (Snijders Blok et al., 2015; Wang et al., 2018); a recent report also suggests that these patients have high rates of neuroblastoma, a rare NC-derived childhood tumor (Lennox et al.). Because this spectrum of non-CNS symptoms is typically caused by impaired NC development (Dubey and Saint-Jeannet, 2017; Vega-Lopez et al., 2018), we hypothesized that *DDX3X* mutations can lead to not only CNS defects but also neurocristopathies.

In this study, we investigated the roles of DDX3 in NC development using *X. tropicalis* as a model. Knockdown (KD) of DDX3 in *X. tropicalis* embryos led to inhibited NC induction and hypoplasia in craniofacial cartilage. Wild-type human DDX3X, but not an RNA helicase-dead mutant, rescued the NC induction defects caused by DDX3 KD. In both *X. tropicalis* embryos and human cells, DDX3 KD reduced endogenous AKT activity and Wnt signaling. The latter could be attributed to decrease in AKT-mediated GSK3β phosphorylation and inhibition, which resulted in enhanced degradation of β-catenin. In addition, Snai1, another GSK3β substrate, was also reduced post-transcriptionally. The defects in NC induction caused by DDX3 KD could be rescued by a constitutively active form of AKT, or by co-expression of β-catenin and Snai1. Finally, DDX3 KD led to decreased levels of RAC1, a known downstream target of DDX3 and upstream activator of AKT, and exogenous RAC1 also rescued the NC induction phenotypes caused by DDX3 KD in *X. tropicalis* embryos. Collectively, our data suggest a critical role of DDX3 in regulating the AKT-GSK3β signaling axis through RAC1, and provide a mechanistic explanation for the similar neurocristopathies that have been observed in patients with mutations in *DDX3X*, *RAC1*, or *AKT*-related genes.

## Results

### DDX3 is required for normal NC induction and craniofacial morphogenesis in *X. tropicalis* embryos

We first performed *in situ* hybridization to determine the expression of *ddx3* during early *Xenopus* development. As reported previously (Cruciat et al., 2013), *ddx3* is expressed ubiquitously in the animal half of the embryos at early gastrula stages (Fig. S1A and S1A’). The expression was primarily found in the dorsal ectoderm at the end of gastrulation (Figs. S1B and S1B’), and in the neural plate with relatively high levels at the NPB during neurulation (Figs. S1C and S1D). At tailbud stages, *ddx3* mRNA was detected in the head and somites (Fig. S1E). Thus, the expression of *ddx3* is consistent with its potential function in CNS and NC development. To understand the roles of DDX3 in NC development and craniofacial morphogenesis, we used a new transgenic *X. tropicalis* line expressing enhanced green fluorescent protein (eGFP) driven by the *snai2* promoter/enhancer. The strong eGFP expression in the differentiating cranial NC lineage allows direct observation of craniofacial cartilage development in live *snai2:eGFP* embryos (Li et al., 2019). Injection of a well-characterized translation-blocking DDX3 morpholino (MO), which targets the 5’-untranslated region (5’-UTR) of the *X. tropicalis ddx3* mRNA (Cruciat et al., 2013), led to hypoplasia of craniofacial cartilage structures on the injected side, as compared to the uninjected side (Fig. 1A). The same phenotype was obtained with a second MO that targets a different site (MO DDX3-2), confirming that this phenotype is specific for DDX3 KD (Fig. 1A). These results suggest that, similar to human DDX3X, *Xenopus* DDX3 is also important for craniofacial morphogenesis.

**Figure 1.**
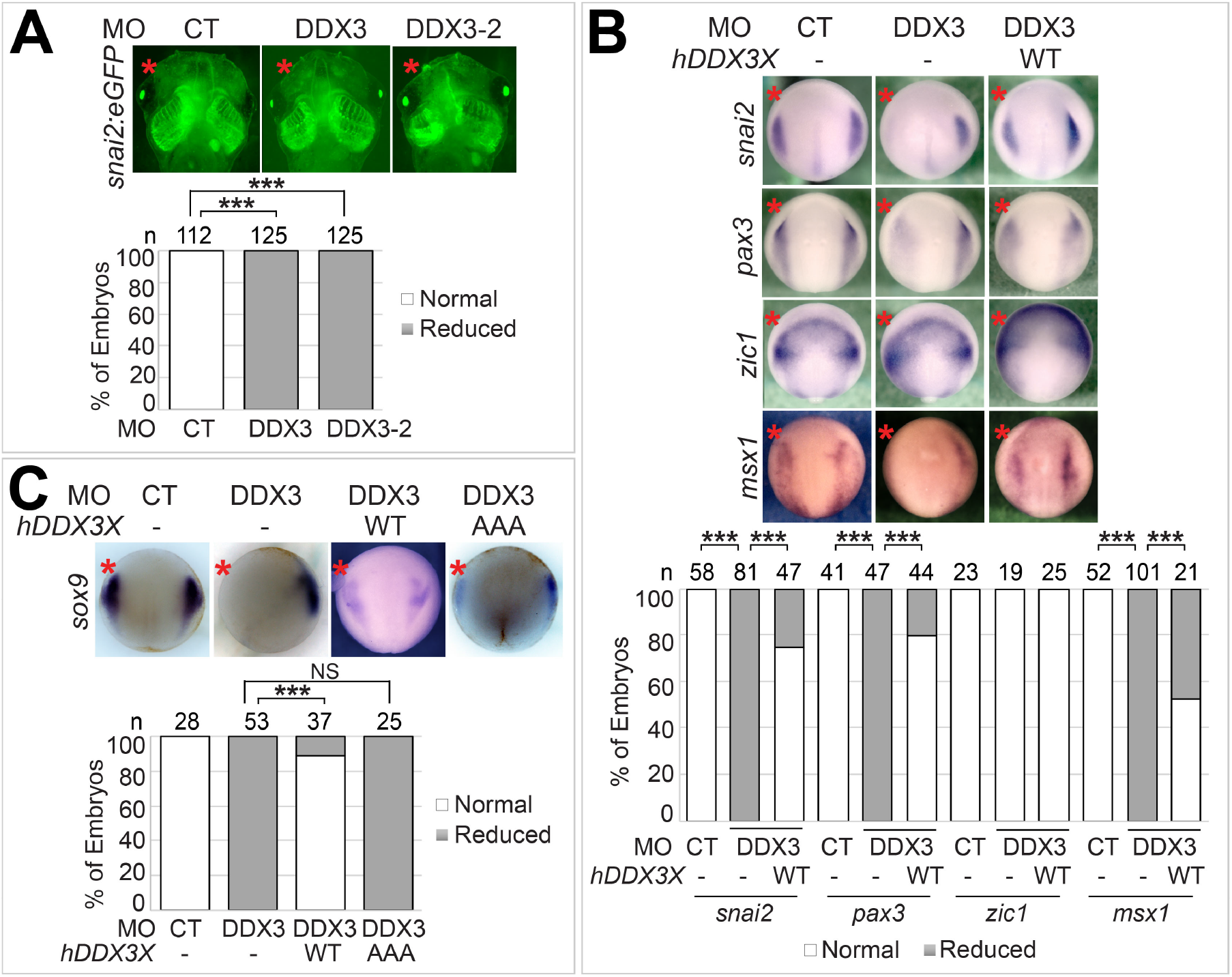
DDX3 is necessary for normal NC induction and craniofacial cartilage development. **A.** One anterodorsal (D1) blastomere of 8-cell stage *snai2:eGFP* embryos was injected with the indicated MO (1.5 ng each). Embryos were cultured to stage ∼46 and imaged for eGFP expression. **B** and **C.** Wild-type *X. tropicalis* embryos were injected in one blastomere at 2-cell stage with the indicated MO (6 ng each) and mRNA encoding wild-type human DDX3X or the AAA mutant (200 pg each), cultured to stage ∼12.5, and processed for *in situ* hybridization for the indicated markers. Representative embryos are shown in ventral (**A**) or dorsal (**B** and **C**) view with anterior at the top, and injected side is denoted with a red star (same below). CT, control; WT, wild-type; n: number of embryos scored (same below). ***, P<0.001; NS, not significant.

To assess the effects of DDX3 KD on early NC development, we examined the expression of NPB and NC specifiers at the end of gastrulation (stage ∼12.5), when the induction of cranial NC is nearly complete. KD of DDX3 reduced the expression of *snai2* and *sox9*, two NC specifiers, indicating that NC induction is inhibited (Figs. 1B and 1C). Interestingly, different NPB specifiers responded differently to DDX3 KD: while both *pax3* and *msx1* were downregulated, *zic1* expression remained largely unchanged and was slightly expanded in some embryos (Fig. 1B). The expansion of *zic1* was probably secondary to the reduction in PAX3, as KD of PAX3 causes a similar effect (Sato et al., 2005). Consistent with these results, injection of a plasmid encoding *X. tropicalis* DDX3 expanded the expression of *snai2* and *msx1* (Fig. S2). Because loss of either Pax3 or Msx1 can block subsequent NC specification (Monsoro-Burq et al., 2005), we conclude that DDX3 depletion inhibits NC induction by interfering with proper NPB formation, although we cannot rule out the possibility that DDX3 also regulates NC specification directly. Co-injection of wild-type human *DDX3X* mRNA restored the expression of all markers to normal levels (Figs. 1B and 1C), further confirming the specificity of the MO and conservation of DDX3 function in NC induction between frogs and humans. Nearly all missense *DDX3X* mutations associated with human birth defects affect evolutionarily conserved residues within the two RNA helicase domains, and some of these mutations have been shown to disrupt the RNA helicase activity of DDX3X (Lennox et al.). We therefore tested if DDX3 RNA helicase activity is needed for NC induction. As shown in Fig. 1C, the S382A/T384A (“AAA”) mutant of human DDX3X, which cannot unwind RNA but retains the ATPase activity (Chao et al., 2006), did not rescue *sox9* expression in DDX3 morphants, suggesting that the RNA helicase activity of DDX3 is important for its function in NC induction.

### DDX3 can regulate β-catenin levels through an RNA helicase-dependent mechanism

DDX3 is known to be a key regulator of Wnt/β-catenin signaling (Cruciat et al., 2013), and the differential responses of NPB specifiers to DDX3 KD (Fig. 1B) are consistent with previous reports showing that Wnt signaling induces the expression of *pax3* and *msx1* but not *zic1* (de Croze et al., 2011; Hong and Saint-Jeannet, 2007; Li et al., 2009). To investigate directly the roles of DDX3 in Wnt signaling during NC induction, we took advantage of a transgenic Wnt reporter line (Tran et al., 2010). Using this reporter line, we showed recently that there is strong endogenous Wnt signaling at the NPB during NC induction (Li et al., 2018). As expected, this Wnt activity was inhibited by KD of DDX3 (Fig. 2A). The transcription factor GBX2 is a direct Wnt target that mediates β-catenin-induced *pax3* and *msx1* expression during *Xenopus* NC induction (Li et al., 2009). Similarly, when human ES and iPS cells are induced to differentiate into NC cells, *gbx2* is expressed prior to the NPB specifiers such as *pax3* and *msx1* in a β-catenin-dependent manner (Leung et al., 2016). In embryos injected with the DDX3 MO (“morphants”), there was a reduction of *gbx2* at stage ∼12 that can be rescued by the mRNA encoding wild-type human DDX3X (Fig. 2B). Conversely, ectopic DDX3 expanded the expression of *gbx2* (Fig. S2), further supporting an important role for DDX3 in Wnt signaling in NC induction.

**Figure 2.**
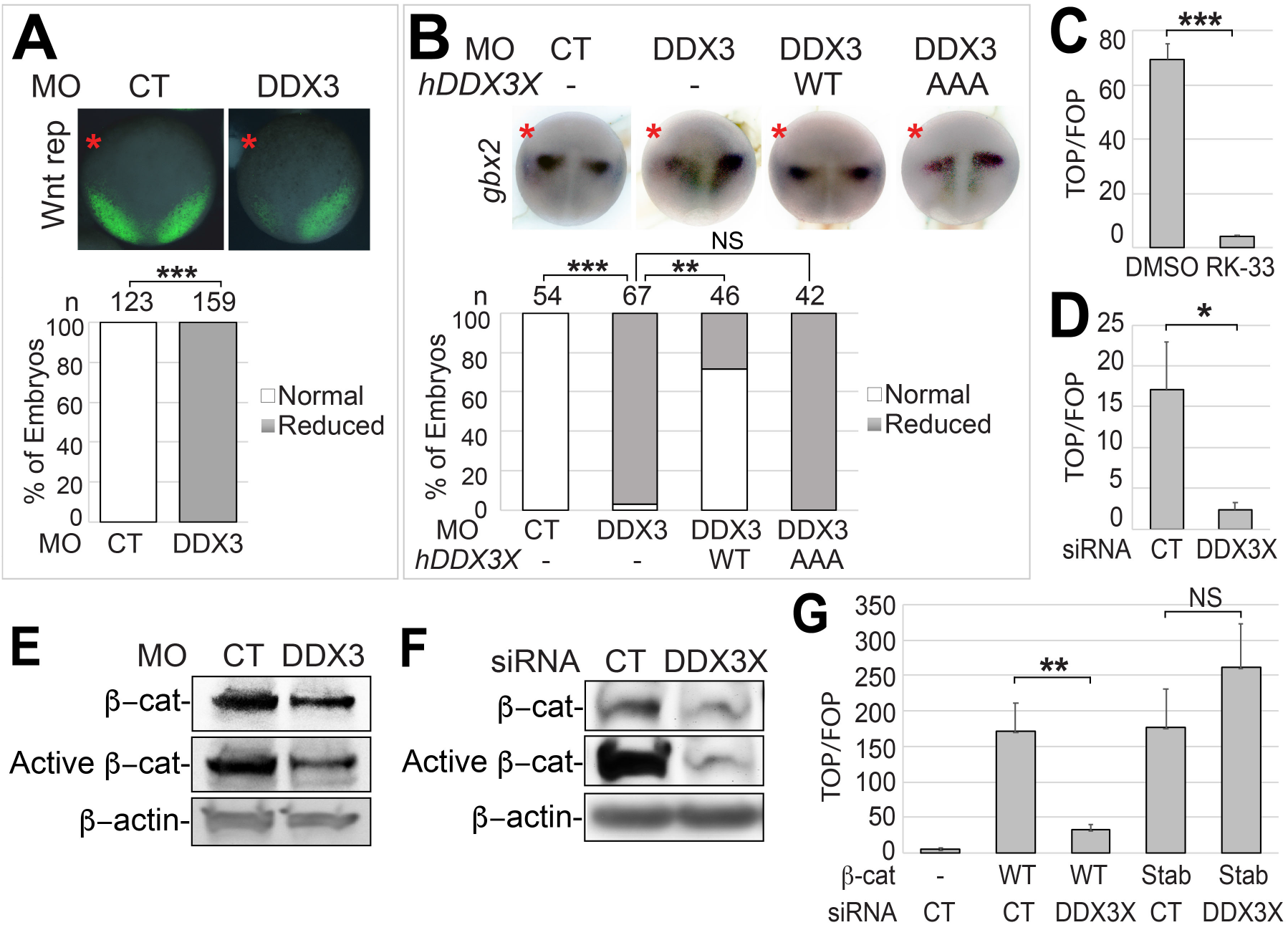
DDX3 stabilizes β-catenin and activates Wnt signaling. Wnt reporter (**A**) or wild-type (**B**) embryos were injected in one blastomere at 2-cell stage with the indicated MO (6 ng each) and mRNA (200 pg), cultured to stage ∼12.5 (**A**) or ∼12 (**B**), and imaged for eGFP expression (**A**) or processed for *in situ* hybridization for *gbx2* (**B**). Representative embryos are shown in dorsal view with anterior at the top. **C**, **D**, **F** and **G**. HEK293T cells were treated with DMSO or 10 μM RK-33 for 24 hr (**C**), or transfected with the indicated siRNA (200 nM) and a plasmid encoding wild-type β-catenin or the stabilized mutant (Stab; 1 μg per well for 6-well plates) for 48 hr (**D**, **F** and **G**). Cell lysates were processed for TOP/FOPFLASH assays (**C**, **D** and **G**), or western blot analyses using the indicated antibodies (**F**). **E.** One-cell stage *X. tropicalis* embryos were injected with the indicated MO (12 ng each) and cultured to stage ∼12.5. Embryo lysates were processed for western blot analyses using the indicated antibodies. *, P<0.05; **, P<0.01; ***, P<0.001; NS, not significant.

Activation of the Wnt signaling pathway starts with binding of the soluble Wnt ligands to cell-surface Frizzled receptors and LRP5/6 co-receptors, which leads to inhibition of glycogen synthase kinase 3 beta (GSK3β)-mediated β-catenin phosphorylation and degradation. Consequently, β-catenin accumulates and enters the nucleus, where it interacts with LEF/TCF family transcription factors to control target gene expression (Loh et al., 2016; Nusse and Clevers, 2017). DDX3 has been shown to regulate Wnt signaling by stimulating the casein kinase CK1ε to phosphorylate the adaptor protein Dishevelled, resulting in enhanced β-catenin nuclear import. This mechanism is independent of the RNA helicase activity of DDX3, as mutants with the ATP-binding domain and part of the helicase C-terminal domain deleted can still activate Wnt signaling (Cruciat et al., 2013). However, unlike wild-type human DDX3X, the AAA mutant had reduced activity in rescuing *gbx2* expression in DDX3 morphants (Fig. 2B), suggesting that DDX3 RNA helicase activity is involved in Wnt signaling. This is consistent with previous studies showing that RK-33, a selective DDX3 RNA helicase inhibitor, inhibits Wnt signaling in multiple types of cancer cells (Heerma van Voss et al., 2017; Heerma van Voss et al., 2015; Tantravedi et al., 2018). Because cancer cells usually harbor many mutations, which could interfere with the action of DDX3, we tested the effects of RK-33 in the non-tumorous HEK293T cells. As in cancer cells, RK-33 also inhibits endogenous Wnt signaling in HEK293T cells, and a similar effect was observed with a DDX3X siRNA (Figs. 2C and 2D). Together, these results point to an important role of DDX3 RNA helicase activity in Wnt signaling and NC induction.

We next examined if DDX3 can regulate Wnt signaling via a mechanism that is different from what was described previously. Interestingly, in both *X. tropicalis* DDX3 morphants and HEK293T cells treated with DDX3X siRNA, we detected decreased levels of total and active (unphosphorylated) β-catenin (Figs. 2E and 2F), which cannot be explained by the helicase-independent mechanism that affects β-catenin nuclear import only (Cruciat et al., 2013). To test if DDX3 function in Wnt signaling is dependent on β-catenin degradation, we compared the effects of DDX3X siRNA on Wnt signal activities induced by wild-type β-catenin and a stabilized mutant that cannot be phosphorylated by GSK3β. As shown in Fig. 2G, KD of DDX3X effectively reduced the Wnt activity induced by wild-type β-catenin but not by the stabilized mutant, suggesting that DDX3 can activate Wnt signaling via stabilizing β-catenin.

### Depletion of DDX3 leads to reduced AKT activity and AKT-dependent GSK3β phosphorylation

The stability of β-catenin is mainly controlled by GSK3β-mediated phosphorylation, and GSK3β activity can be regulated by upstream Wnt ligand-receptor interaction or other signal inputs. Among the latter, the serine/threonine kinase AKT can inhibit GSK3β activity via direct phosphorylation of the Ser9 residue (Cross et al., 1995; Fukumoto et al., 2001; Naito et al., 2005; Sharma et al., 2002). Mutations that cause gain of AKT activity in humans and mice lead to overgrowth phenotypes in the CNS and NC (Akgumus et al., 2017; Butler et al., 2005; Rivière et al., 2012), which are opposite to those caused by loss of DDX3X function (Fig. 1). In addition, phosphatidylinositol 3-kinase (PI3K), a kinase that activates AKT, regulates NC induction in *Xenopus laevis*, a species closely related to *X. tropicalis*, although the exact roles of PI3K in NC induction remain controversial (Geary and LaBonne, 2018; Pegoraro et al., 2015). We therefore asked if DDX3 KD interferes with the AKT-GSK3β signaling axis. AKT is activated through two consecutive phosphorylation events, the first occurring at Thr308 and the second at Ser473 (Manning and Toker, 2017). In HEK293T cells, siRNA-mediated KD of DDX3X inhibited both phosphorylation events without affecting total AKT levels (Fig. 3A). Similarly, MO-mediated DDX3 KD also blocked Ser473 phosphorylation in *X. tropicalis* (Fig. 3B); our pThr308 antibody did not pick up the endogenous pThr308 signal in frog embryos. Consistent with the reduced AKT activity, the phosphorylation of GSK3β at Ser9, but not the total GSK3β levels, was also reduced upon DDX3X KD or treatment with RK-33 in HEK293T cells (Figs. 3C and 3D). Because phosphorylation at Ser9 inhibits GSK3β activity, these results provide an explanation for the decrease in β-catenin level when DDX3 activity was reduced (Figs. 2E and 2F). Besides β-catenin, the transcription factor Snai1 is also phosphorylated by GSK3β and subsequently targeted to degradation (Yook et al., 2005; Zhou et al., 2004). Ectopically expressed Snai1 was clearly reduced when DDX3 was knocked down in *X. tropicalis* embryos or HEK293T cells (Figs. 3E and 3F), indicating that DDX3 upregulates Snai1 protein levels post-transcriptionally. Together, our data suggest that DDX3 is required for AKT activation and downstream GSK3β inhibition in *Xenopus* embryos and human cells.

**Figure 3.**
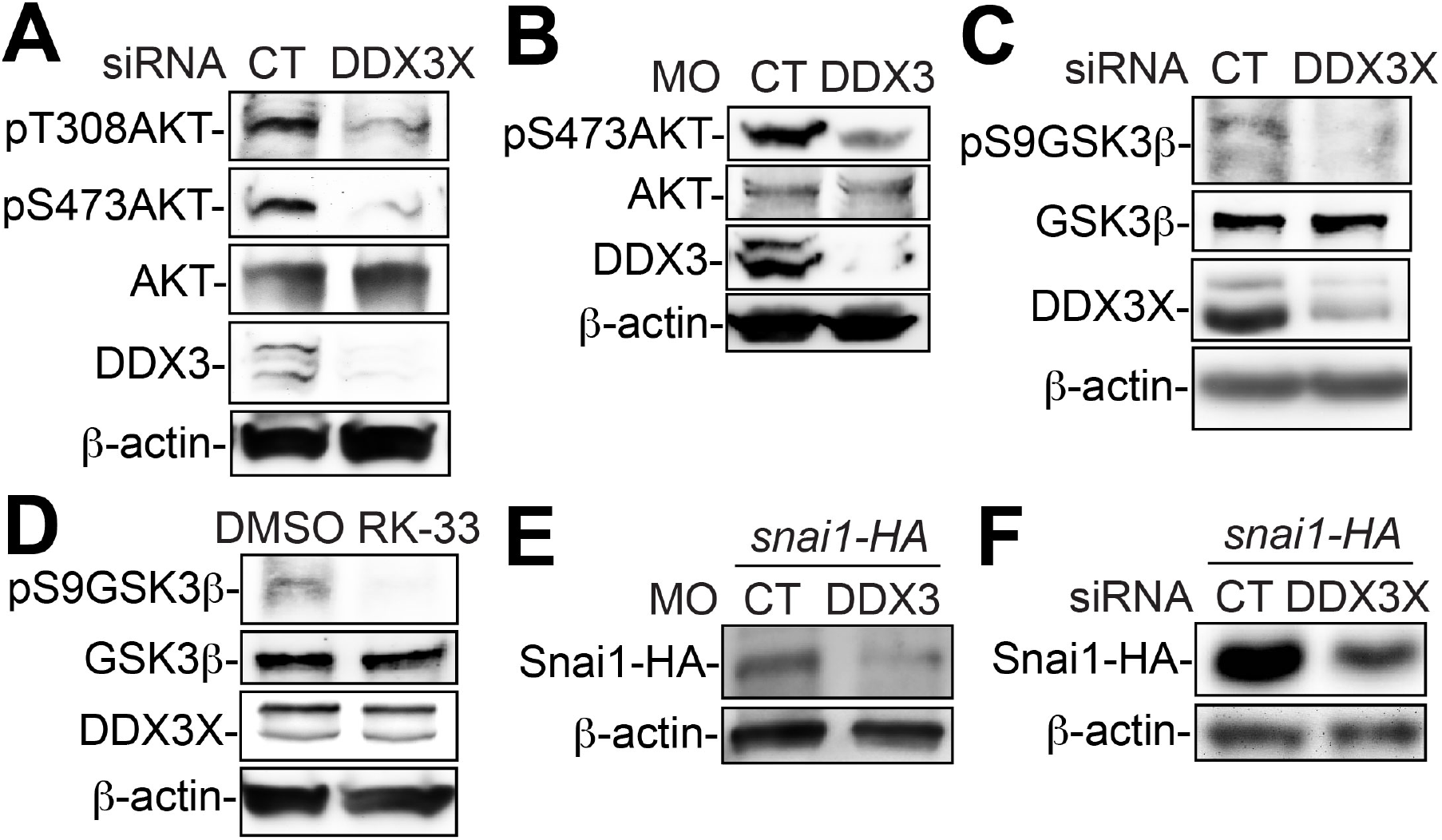
KD of DDX3 reduces AKT activity, AKT-mediated GSK3β phosphorylation and ectopically expressed Snai1. **A**, **C**, **D** and **F**. HEK293T cells were treated with DMSO or 10 μM RK-33 for 24 hr (**D**), or transfected with the indicated siRNA (200 nM) and a plasmid encoding HA-tagged Snai1 (1 μg per well for 6-well plates) for 48 hr (**A**, **C** and **F**). **B** and **E**. Embryos were injected at one-cell stage with the indicated MO (12 ng each) and mRNA encoding Snai1 (50 pg), and cultured to stage ∼12.5. Cell or embryo lysates were processed for western blot analyses using the indicated antibodies.

### AKT is a key regulator of Wnt/β-catenin signaling and NC induction

The function of PI3K-AKT signaling in NC induction is controversial. While treatment of gastrula-stage *X. laevis* embryos with a PI3K inhibitor reduces the expression of NC markers, a constitutively active form of PI3K can also reduce NC markers induced by Pax3 and Zic1 in isolated *X. laevis* animal caps, which are thought to contain pluripotent stem cells (Buitrago-Delgado et al., 2015; Geary and LaBonne, 2018; Pegoraro et al., 2015). However, PI3K has AKT-independent function (Faes and Dormond, 2015), and the mechanisms underlying the possible roles of AKT in NC induction remains largely unknown. To address these questions, we first determined the patterns of NPB and NC markers in embryos treated with AKT inhibitor IV (AKTi), which blocks AKT activity downstream of PI3K. Because inhibition of AKT activity at earlier stages caused gastrulation defects, we cultured the embryos in AKTi from stage ∼10 to ∼12.5. This transient inhibition of AKT activity reduced the expression of NC specifiers *snai2* and *sox9* as well as the NPB specifier *msx1*, but not the other NPB specifier *zic1* (Fig. 4A). These effects are similar to those caused by DDX3 KD (Figs. 1B and 1C). As discussed above, *msx1* but not *zic1* is induced by Wnt signaling, and AKT can stabilize β-catenin by directly inhibiting GSK3β activity. We therefore hypothesized that AKT is required for activating the Wnt signaling pathway, one of the most important signaling pathways in NC induction. To test this hypothesis directly, we treated the Wnt reporter embryos with AKTi, and indeed observed a reduction in the endogenous Wnt activity at the NPB (Fig. 4B). A similar reduction was obtained with the injection of an mRNA encoding a dominant-negative AKT mutant (dnAKT; Fig. 4C), confirming that AKT is a key regulator of Wnt signaling during NC induction.

**Figure 4.**
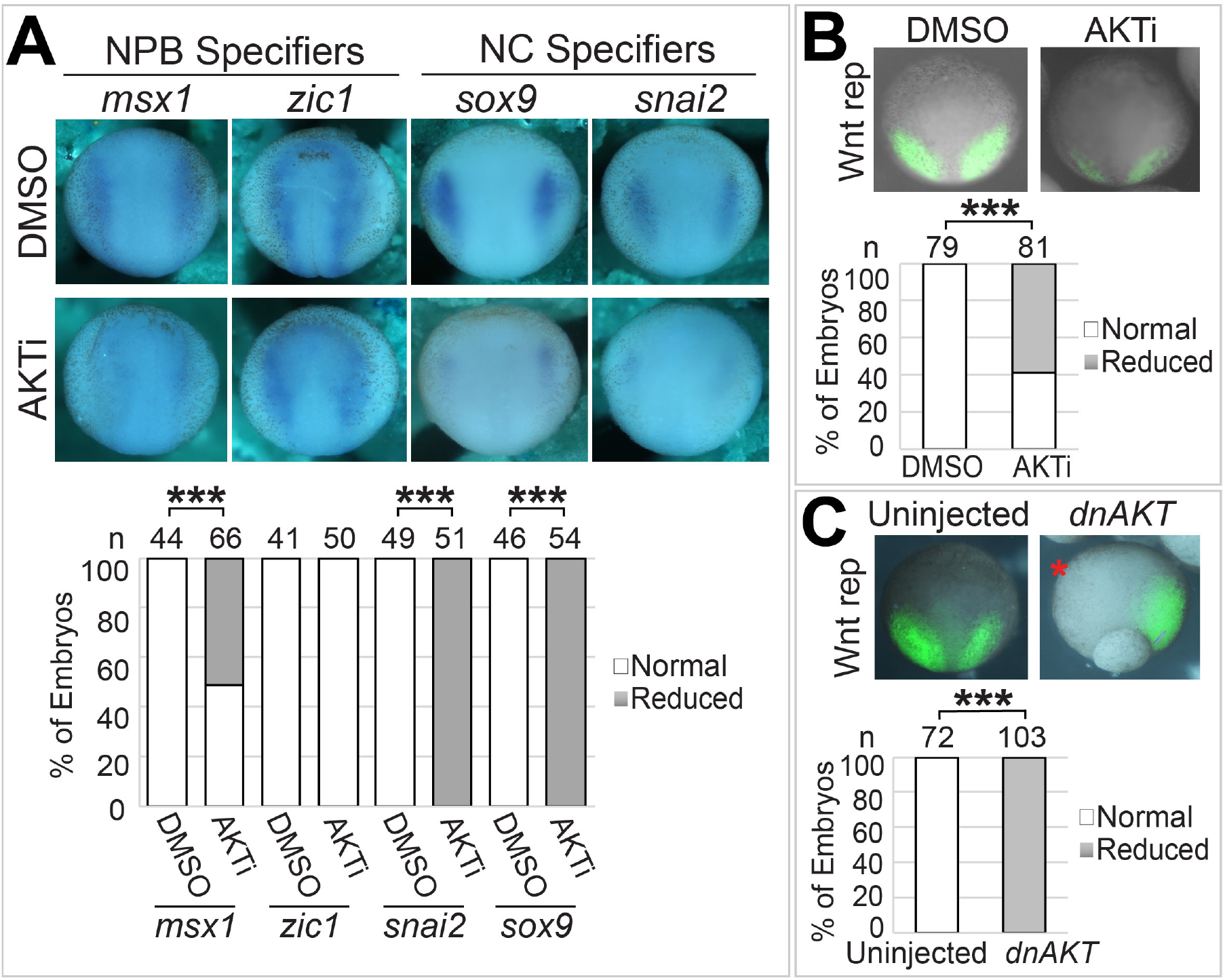
AKT activity is required for Wnt signaling and NC induction. **A** and **B.** Wild-type (**A**) or Wnt reporter (**B**) embryos were treated with 20 μM AKTi or DMSO from stage ∼10 to ∼12.5, and processed for *in situ* hybridization for the indicated markers (**A**) or imaged for eGFP expression (**B**). **C.** Wnt reporter embryos were injected in one blastomere at 2-cell stage with *dnAKT* mRNA (50 pg), cultured to stage ∼12.5, and imaged for eGFP expression. Representative embryos are shown in dorsal view with anterior at the top. ***, P<0.001.

### DDX3 induces the NC by regulating AKT and the GSK3β substrates β-catenin and Snai1

To establish the relationship between DDX3 and its downstream targets in NC induction, we carried out a series of rescue experiments. AKT is recruited to the plasma membrane, where it is activated, through the interaction of its pleckstrin homology (PH) domain and the phosphorylated lipid products of PI3K (Manning and Toker, 2017). Replacement of the PH domain by a myristoylation signal results in a constitutively active form of AKT (caAKT) that functions independently of PI3K (Kohn et al., 1996). We found that overexpression of caAKT induced Wnt activity in HEK293T cells, and this activity was not affected by DDX3X KD (Fig. S3). This is in line with our hypothesis that AKT functions downstream of DDX3 to activate Wnt signaling. We tested further if caAKT can rescue the phenotypes caused by DDX3 KD in *X. tropicalis* embryos. Indeed, co-injection of the *caAKT* mRNA restored *snai2* expression and Wnt signaling at the NPB in DDX3 morphants (Figs. 5A and 5B), confirming the roles of AKT in mediating DDX3 function in Wnt signaling and NC induction. We have shown that KD of DDX3 reduced the inhibitory phosphorylation of GSK3β at Ser9, and have identified β-catenin and Snai1 as two GSK3β substrates whose levels are regulated by DDX3 (Figs. 2E, 2F and 3C-3F). Because both of these GSK3β substrates are essential for NC induction in *Xenopus* (Aybar et al., 2003; LaBonne and Bronner-Fraser, 1998), we tested if they are downstream effectors of DDX3 in NC induction. Ectopic expression of either β-catenin or Snai1 could partially rescue the NPB specifier *msx1* and the NC specifier *sox9*, and a combination of both GSK3β substrates resulted in a nearly complete rescue (Fig. 5C). Based on these results, we conclude that DDX3 functions in NC induction by regulating AKT as well as β-catenin and Snai1.

**Figure 5.**
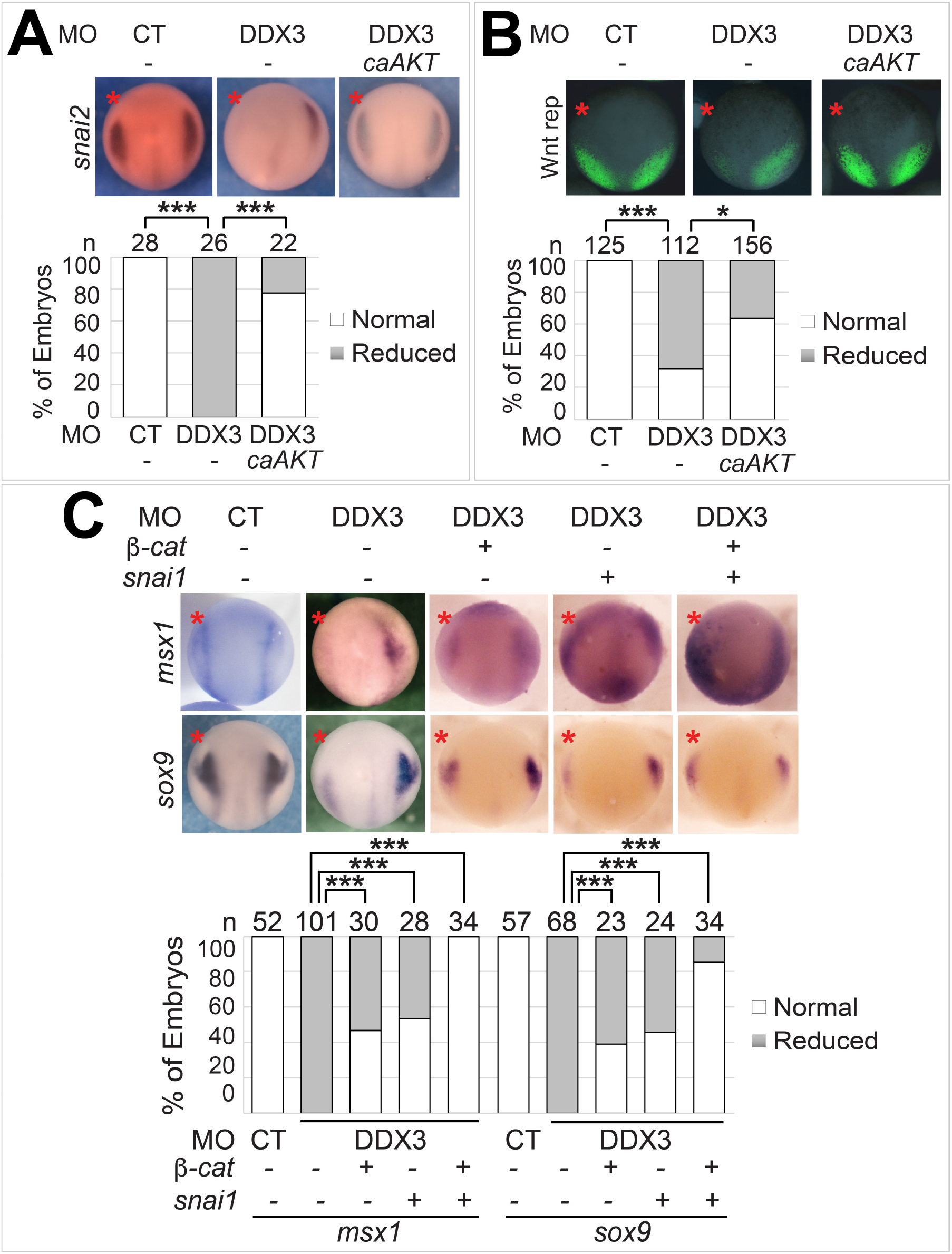
DDX3 induces the NC through AKT, β-catenin, and Snai1. Wild-type (**A** and **C**) or Wnt reporter (**B**) embryos were injected in one blastomere at 2-cell stage with the indicated MO (6 ng each), mRNA (50 pg for caAKT and 100 pg for Snai1), and plasmid (10 pg for *β-catenin*), cultured to stage ∼12.5, and processed for *in situ* hybridization for the indicated markers (**A** and **C**) or imaged for eGFP expression (**B**). Representative embryos are shown in dorsal view with anterior at the top. *, P<0.05; ***, P<0.001.

### RAC1 is likely an immediate downstream effector of DDX3 in NC induction

We next asked how DDX3 activates AKT and the downstream signaling events. DDX3 can promote translational initiation of mRNAs with long and structured 5’-UTR through its RNA helicase activity, a function that is conserved from yeast to humans (Guenther et al., 2018; Sen et al., 2015). One of these mRNAs encodes RAC1, which can mediate DDX3 function in other physiological processes in mammals (Chen et al., 2015; Chen et al., 2016; Ku et al., 2018). Notably, RAC1 can activate AKT through its downstream effector p21-activated kinase (PAK) (Higuchi et al., 2008). In *Xenopus* embryos, a dominant-negative mutant of RAC1 reduces, whereas a constitutively active mutant expands, the expression of NC specifiers (Broders-Bondon et al., 2007). We therefore examined if RAC1 functions downstream of DDX3. To do this, we obtained an *X. tropicalis rac1* cDNA with a 257-nucleotide 5’-UTR. Although the 5’-UTR of *X. tropicalis rac1* did not show any significant sequence similarity to that of human *RAC1*, prediction using the Mfold algorithm suggests that they contain similar secondary structures that are energetically stable (Fig. S4). Thus, the mechanism of translational regulation of RAC1 may be conserved. As shown in Figs. 6A-6C, blocking DDX3 activity by RK-33 in HEK293T cells, or KD of DDX3 in either *X. tropicalis* embryos or HEK293T cells, resulted in a decrease in endogenous RAC1 protein, indicating that RAC1 is a target of DDX3 RNA helicase activity in both frogs and humans. Further, an *X. tropicalis rac1* mRNA without the 5’-UTR rescued the craniofacial defects caused by DDX3 KD (Fig. 6D), suggesting that RAC1 mediates DDX3 function in NC development.

**Figure 6.**
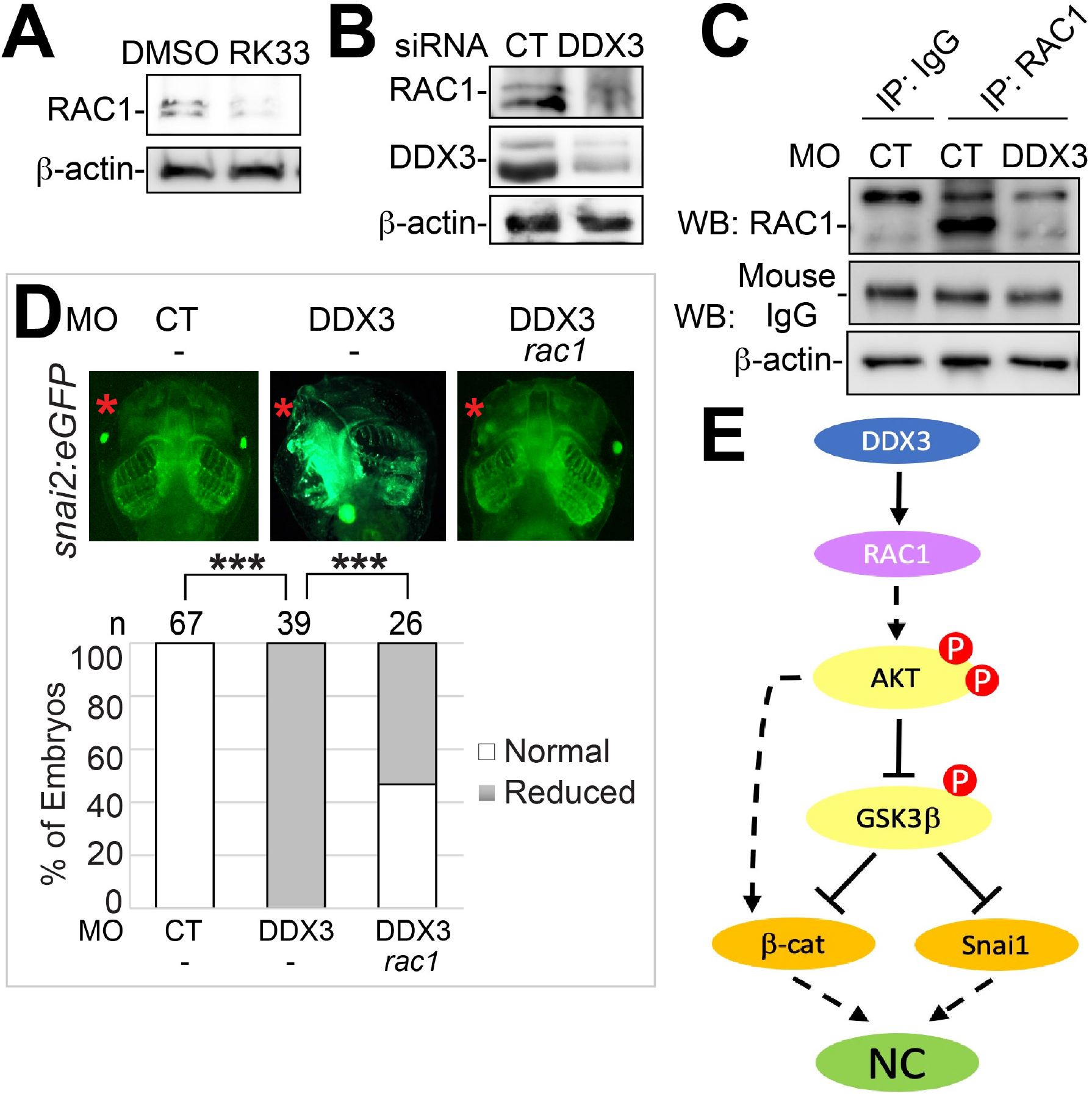
RAC1 is a downstream effector of DDX3 in NC induction. **A** and **B**. HEK293T cells were treated (**A**) or transfected (**B**) as in Fig. 3. Cell lysates were processed for western blotting with the indicated antibodies. **C.** Wild-type embryos were injected at one-cell stage with the indicated MO, and cultured to stage ∼12.5. To obtain a clean signal from the embryo lysates, two RAC1 antibodies were used, one for immunoprecipitation (IP) and the other for western blot (WB) detection (see *Materials and Methods*). Whole-cell lysates were also processed for western blot for β-actin. **D.** One anterodorsal (D1) blastomere of 8-cell stage *snai2:eGFP* embryos was injected with the indicated MO (1.5 ng each) and mRNA (100 pg). Embryos were cultured to stage ∼46 and imaged for eGFP expression. Representative embryos are shown in ventral view with anterior at the top. **E.** A model for DDX3 function in regulating downstream signaling during NC induction.

## Discussion

First discovered in 2015, mutations in DDX3 have drawn a great deal of research interest as a frequent cause of intellectual disability in humans (Dikow et al., 2017; Kellaris et al., 2018; Lennox et al.; Nicola et al., 2019; Snijders Blok et al., 2015; Wang et al., 2018). Although nearly all published studies on patients with DDX3 mutations are focused on the CNS defects, we noted that the majority of these patients also have abnormalities in craniofacial structures and/or other NC-derived tissues, which have not been addressed previously. We therefore hypothesized that DDX3 plays important roles in NC development, and provide here the first evidence supporting this hypothesis. Based on our data, we propose that the RNA helicase activity of DDX3 is required for efficient translation of RAC1, which activates AKT to phosphorylate and inhibit GSK3β. The inhibition of GSK3β leads to stabilization of β-catenin and Snai1, two proteins essential for NC induction (Fig. 6E).

While DDX3 has been shown to activate Wnt signaling through an RNA helicase-independent mechanism, several lines of evidence suggest the existence of an additional helicase-dependent mechanism. Missense *DDX3X* mutations associated with human birth defects cause decrease in RNA helicase activity, and assays with zebrafish embryos indicate that these mutations impair DDX3X’s ability to activate Wnt signaling (Lennox et al.; Snijders Blok et al., 2015). Further, RK-33, a selective inhibitor of DDX3 RNA helicase activity, reduces Wnt signaling in several cancer cell lines (Bol et al., 2015; Tantravedi et al., 2018). Similarly, RK-33 inhibits Wnt signaling in the non-tumorous HEK293T cells in our reporter assays, and the helicase-dead AAA mutant did not rescue the reduced expression of the Wnt target gene *gbx2* in DDX3 morphants (Fig. 2B). In addition, we uncovered an effect of DDX3 KD on β-catenin levels, which is consistent with the downregulation of AKT activity instead of the helicase-independent mechanism described previously (Cruciat et al., 2013). Thus, our data suggest that DDX3 can also regulate Wnt signaling through its RNA helicase activity.

The RNA helicase activity of DDX3 is needed for the efficient translation of certain mRNAs with highly structured 5’-UTR (Guenther et al., 2018; Sen et al., 2015). A known direct target of DDX3 in mammals is RAC1 (Chen et al., 2015; Chen et al., 2016; Ku et al., 2018), which is extremely conserved throughout vertebrate evolution (100% identical between human and *X. tropicalis* orthologs). Interestingly, mutations in *RAC1* cause craniofacial disorders and other potential NC and CNS defects, which are highly similar to those caused by *DDX3X* mutations (Reijnders et al., 2017). In line with these observations, *rac1* is expressed in the NC and CNS in *Xenopus* embryos, and has been implicated in *Xenopus* NC induction (Broders-Bondon et al., 2007; Lucas et al., 2002). KD of DDX3 leads to reduced RAC1 protein levels in *X. tropicalis* embryos, and exogenous RAC1 rescues the craniofacial defects caused by DDX3 KD (Figs. 6C and 6D), suggesting that RAC1 mediates DDX3 function in NC induction. RAC1 can activate AKT through its immediate downstream effector PAK (Higuchi et al., 2008). Among the vertebrate PAK family members, four (PAK1-4) contain a CDC42/RAC interaction/binding (CRIB) motif within the N-terminal regulatory domain, which binds and inhibits the C-terminal kinase domain when PAK is inactive. The direct interaction of the CRIB motif with activated RAC1 leads to dissociation of the regulatory domain from the kinase domain and activation of PAK (Zhao and Manser, 2012). The freed kinase domain of PAK1 can serve as a scaffold to bind AKT to facilitate its membrane localization and activation through a kinase-independent mechanism (Higuchi et al., 2008), but a separate study suggests that PAK1 activates AKT by phosphorylating AKT directly at Ser473 (Mao et al., 2008). Hence it would be of interest to test whether PAK also functions downstream of DDX3 in NC induction and, if so, this function is dependent on PAK kinase activity.

Previous studies suggest that PI3K, an upstream activator of AKT, can either promote or inhibit NC induction (Geary and LaBonne, 2018; Pegoraro et al., 2015). Here we show that direct inhibition of AKT activity reduces endogenous Wnt signaling at the NPB, causing altered expression of NPB and NC markers that resembles loss of DDX3 or Wnt signaling (Figs. 4A-4C). These data indicate that one mechanism through which AKT functions in NC induction is by regulating Wnt signaling, a major signaling pathway that is critical for NC induction. It should be noted, though, that our results do not contradict with the model that excessive PI3K/AKT activities inhibit NC induction. In fact, we found that high levels of ectopic DDX3 can cause reduction in NPB and NC markers (data not shown), suggesting that stringently controlled AKT activity is required for proper NC induction.

AKT-mediated phosphorylation of GSK3β at Ser9 inhibits GSK3β activity, leading to the stabilization of certain GSK3β substrates (Cross et al., 1995; Zhou et al., 2004). However, although some studies show that AKT promotes Wnt signaling, likely by inhibiting GSK3β and stabilizing β-catenin (Fukumoto et al., 2001; Naito et al., 2005; Sharma et al., 2002), others suggest that AKT and Wnt act on two separate pools of GSK3β and do not crosstalk with each other (Ding et al., 2000; Ng et al., 2009). Besides inhibiting GSK3β, AKT has also been shown to activate Wnt signaling through other mechanisms. For example, PI3K/AKT signaling can upregulate the transcriptionally active β-catenin (containing unphosphorylated Ser37 and Thr41) in HEK293T and other cell lines, possibly by inducing the phosphatase PP2A to dephosphorylate these two residues (Persad et al., 2016). Indeed, we detected a reduction in active β-catenin that was more drastic than in total protein (Figs. 2E and 2F). Thus, AKT may mediate DDX3-induced Wnt signaling through GSK3β-dependent and -independent mechanisms.

Unlike β-catenin, Snai1, another substrate of GSK3β, is generally believed to be stabilized upon AKT-mediated phosphorylation of GSK3β (Zhou et al., 2004). Here we show that KD of DDX3 results in decreased levels of ectopically expressed Snai1 (Figs. 3E and 3F), and that ectopic Snai1 and β-catenin cooperatively rescue the NC induction defects caused by DDX3 KD in *X. tropicalis* embryos (Fig. 5C). These data suggest that DDX3-regulated AKT-GSK3β signaling axis induces NC through Snai1 in addition to β-catenin. Currently, most protocols for inducing the NC from human pluripotent stem cells include GSK3β inhibition, which is thought to activate canonical Wnt signaling through β-catenin stabilization (Gomez et al., 2019; Leung et al., 2016; Menendez et al., 2011; Mica et al., 2013). In light of our results, it would be of interest to examine if the stabilization of Snai1 also contributes to the NC induction abilities of GSK3β inhibitors.

Consistent with our model that DDX3, RAC1, AKT and β-catenin function in the same signaling cascade to regulate NC induction (Fig. 6E), mutations in *RAC1*, *CTNNB1* (encoding β-catenin) and genes that directly control AKT activity (including *AKT1*, *AKT3*, *PIK3R2*, *PIK3CA* and *PTEN*) can all lead to craniofacial disorders and other potential neurocristopathies such as pigment defects, in humans (Akgumus et al., 2017; Butler et al., 2005; Kharbanda et al., 2017; Reijnders et al., 2017; Rivière et al., 2012; Tucci et al., 2014). Current efforts are therefore focused on investigating the effects of human mutations in *DDX3X* and these target genes on downstream signaling and NC induction. In addition, it is worth noting that patients carrying mutations in these genes also display highly similar CNS defects, including intellectual disability, autism spectrum disorder, micro/macrocephaly, and corpus callosum hypo/hyperplasia (Akgumus et al., 2017; Butler et al., 2005; Kharbanda et al., 2017; Reijnders et al., 2017; Rivière et al., 2012; Snijders Blok et al., 2015; Tucci et al., 2014; Wang et al., 2018). Hence it is tempting to test if the signaling cascade that we propose here (Fig. 6E) is also essential for neurogenesis and CNS development. Our study may lay the foundation for future work to understand the pathophysiology of the birth defects caused by mutations that interfere with this signaling cascade.

## Materials and methods

### Plasmids and reagents

Constructs encoding dnAKT (clone 9031), caAKT (clone 10841), FLAG-tagged human β-catenin (clone 16828), and HA-tagged human Snai1 (clone 31697) were purchased from Addgene. Full-length cDNA clones for human *DDX3X* (Accession: BC011819), human *RAC1* (BC050687), and *X. tropicalis gbx2* (Accession: NM_001011472) were from GE-Dharmacon. The cDNAs for human *DDX3X* and *RAC1* were subcloned into a pCS2+ expression vector with an in-frame HA tag, and the S382A/T384A (“AAA”) mutant of DDX3 were generated by PCR using the mutagenesis primers CCACACTATGATGTTTGCTGCTGCTTTTCCTAAGGAAATAC (forward) and GTATTTCCTTAGGAAAAGCAGCAGCAAACATCATAGTGTGG (reverse). Constructs for the expression of *X. laevis* β-catenin and Snai1 and for *in situ* hybridization for *snail2, sox9, foxd3, zic1,* and *msx1* were obtained in previous studies (Li et al., 2018; Wei et al., 2012; Wei et al., 2010). *In vitro* transcription was carried out as described to generate *in situ* hybridization probes and mRNA transcripts (Sive et al., 2000). Pharmacological inhibition of AKT and DDX3X was performed with AKT Inhibitor IV (Calbiochem 124011) and RK-33 (Selleckchem S8246), respectively. Antibodies that were used in this study include mouse anti-myc (DSHB 9E10), mouse anti-HA (Sigma-Aldrich H9658), rabbit anti-DDX3 (Abcam ab151965), rabbit anti-RAC1 (Invitrogen PA5-44839; for western blotting), mouse anti-RAC1 (Abcam ab33186; for immunoprecipitation), rabbit anti-β-catenin (Sigma C2206), rabbit anti-active-β-catenin (Cell Signaling Technology 8814), mouse anti-GSK3β (Santa Cruz sc-53931), rabbit anti-piS9-GSK3β (Invitrogen PA1-4688), rabbit anti-AKT1 (Aviva Systems Biology AVARP06008_P050), rabbit anti-pS473-AKT (Cell Signaling Technology 4060S), rabbit anti-*Xenopus*-pS473-AKT (Cell Signaling Technology 9271S), and rabbit anti-pT308-AKT (Cell Signaling Technology 9275). Secondary antibodies that were used include HRP-conjugated rabbit anti-mouse (Sigma-Aldrich A9044) and goat anti-rabbit (Sigma-Aldrich A0545). Mouse anti-β-actin (Sigma-Aldrich A5316) was used for loading controls.

### Animals and embryo manipulation

Wild-type *X. tropicalis* frogs were purchased from NASCO. The transgenic *X. tropicalis* Wnt reporter frogs were courtesy of Dr. Kris Vleminckx (Flanders Institute for Biotechnology, Belgium), and the transgenic *X. tropicalis snai2:eGFP* line was generated as described (Li et al., 2019). Methods involving live animals were carried out in accordance with the guidelines and regulations approved and enforced by the Institutional Animal Care and Use Committees at West Virginia University and University of Delaware. Embryo were collected and injected with PLI-100A microinjectors (Harvard Apparatus) as described previously (Li et al., 2018). Morpholinos DDX3 (5’-CCGTGATTGATGCTT-3’) and DDX3-2 (5’-TTTCCACGGCCACATGACTCATAAC-3’) were designed and generated by Gene Tools. Alexa Fluor 555 (Invitrogen; for direct phenotype observation) or 488 (Invitrogen; for in situ hybridization) was co-injected as a lineage tracer. Injected embryos were sorted by the co-injected lineage tracer and cultured in 0.1x MBS to desired stages, when the embryos were imaged directly or processed for *in situ* hybridization or western blotting. Bright-field and fluorescence imaging was carried out using a Zeiss Axiozoom v16 epifluorescence microscope, and images of embryos were taken with an AxioCam MRc Rev3 camera. For western blotting of embryo lysates, embryos were lysed as described (Li et al., 2018).

### Cell culture and transfection

HEK293T cells (ATCC), recently authenticated and tested for contamination, were cultured in DMEM (ATCC) supplemented with 10% fetal bovine serum (Gibco) at 37°C with 5% CO_2_. Cells were transfected with DDX3 or control siRNA (Dharmacon J-006874-06-0002 and D-001810-01-05) at 50% confluency or with plasmids at 70% confluency, using Lipofectamine 3000 (Invitrogen). For experiments that used both siRNA and plasmid, siRNA was transfected first, and the media was changed and the plasmid transfected on the following day. Cell lysates were prepared as described previously (Li et al., 2018). For TOP/FOPFLASH assays, cell were transfected and luciferase assays were carried out as described (Wei et al., 2010).

### *In situ* hybridization, western blot and immunoprecipitation

Embryos collected at desired stages were fixed in 4% paraformaldehyde for 24 hours at 4°C, and *in situ* hybridization was conducted subsequently as described (Sive et al., 2000). Western blotting and immunoprecipitation were performed as described previously (Li et al., 2018), and detection was carried out using HRP-conjugated antibodies and chemiluminescence substrates (GE Healthcare).

### Phenotype Scoring and Statistics

Embryos were scored by comparing the injected side with the uninjected side of the same embryos. The percentage of normal and reduced phenotypes were calculated for injected embryos obtained from multiple independent experiments, and Chi-squared tests were performed to compare the phenotypes in different treatment groups. For craniofacial phenotypes, injected *snai2-eGFP* embryos were allowed to develop to stage ∼46 and scored for defects in head cartilage structures. Images of head cartilages (eGFP) were taken with a Zeiss Axiozoom.v16 epifluorescence microscope.

## Acknowledgements

We thank Dr. Kris Vleminckx for providing the transgenic Wnt reporter frogs. This work was supported by the National Institute of Health (GM114105 and GM104316 to SW).

## Competing interests

The authors declare no competing interest.

---

**Figure S1.**
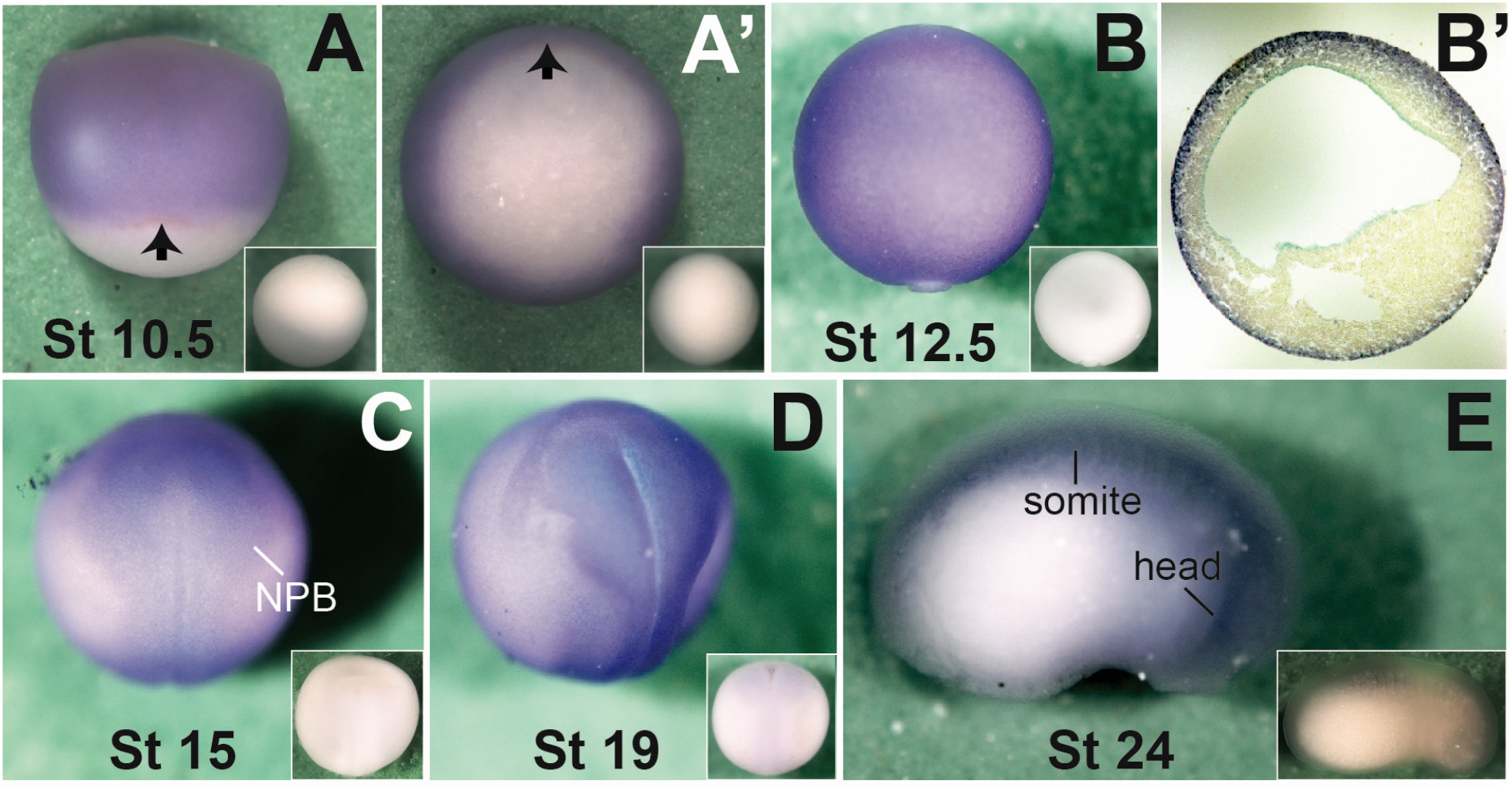
Developmental expression of *ddx3* during *X. tropicalis* embryogenesis. *In situ* hybridization was carried out for *ddx3* with *X. tropicalis* embryos at indicated stages. **A-D**. Dorsal view with animal pole (**A**) or anterior (**B-D**) at the top. **A’.** Vegetal view of the same embryo in **A** with dorsal at the top. **B’.** Transverse section of a stage ∼12.5 embryo with dorsal at the top, showing high *ddx3* expression in the ectoderm. **E.** Side view with anterior to the right and dorsal at the top. Control *in situ* hybridization with sense probe is shown in insets, and arrows in **A** and **A’** indicate the dorsal blastopore lip.

**Figure S2.**
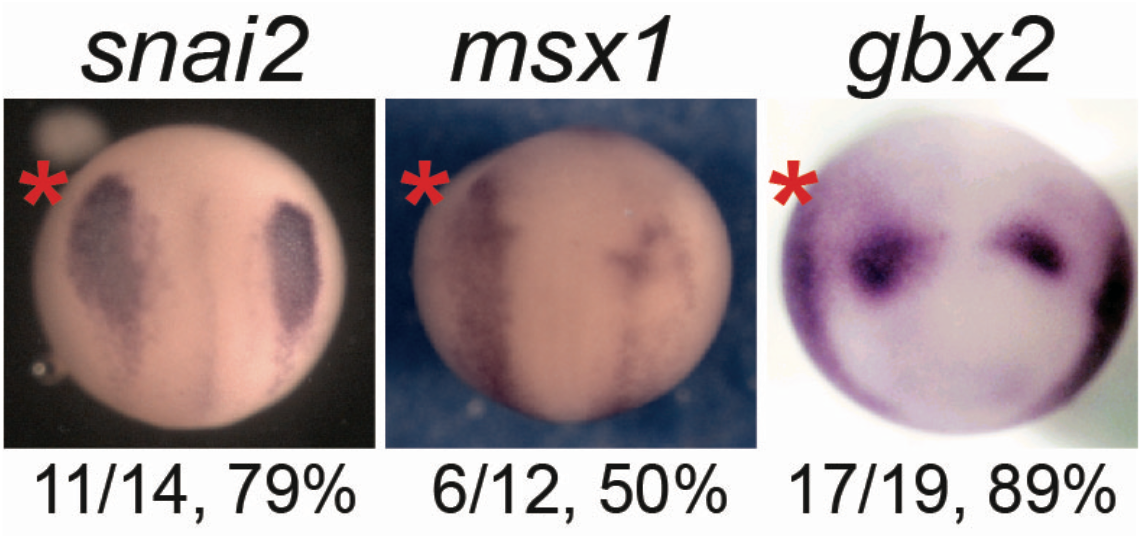
Ectopically expressed *X. tropicalis* DDX3 causes expansion of NPB and NC specifiers. Wild-type *X. tropicalis* embryos were injected in one blastomere at 2-cell stage with 10 pg plasmid encoding *X. tropicalis* DDX3, cultured to stage ∼12.5, and processed for *in situ* hybridization for the indicated markers. Embryos are shown in dorsal view with anterior at the top, with the number and percentage of injected embryos displaying the indicated phenotypes shown underneath.

**Figure S3.**
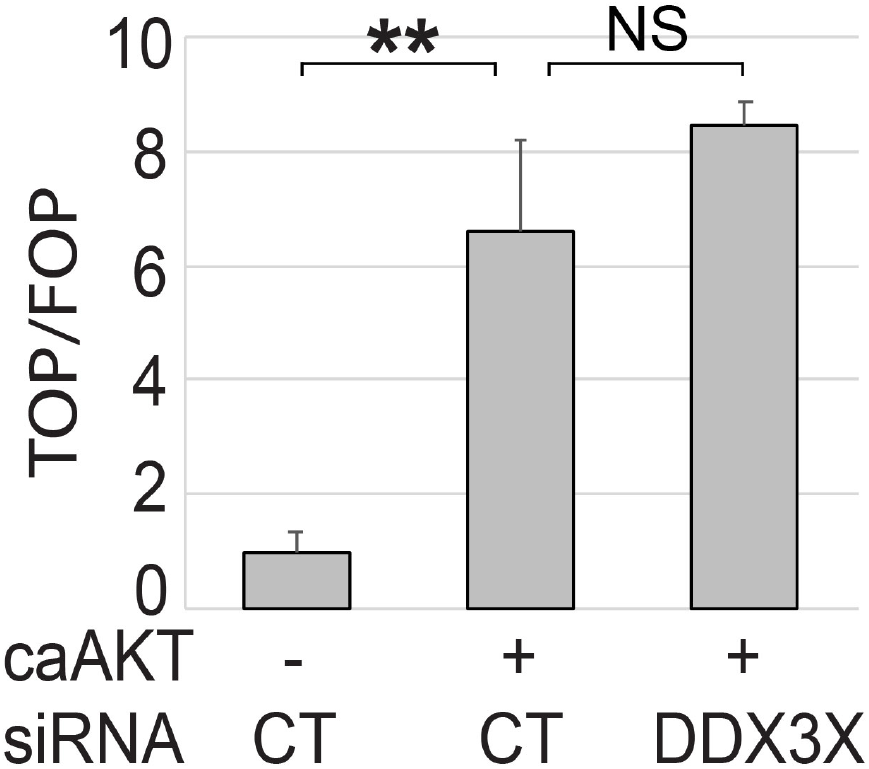
A constitutively active mutant of AKT activates Wnt signaling independently of DDX3. HEK293T cells were transfected with the indicated siRNA (200 nM) and a plasmid encoding caAKT (1 μg per well for 6-well plates) for 48 hr. Cell lysates were processed for TOP/FOPFLASH assays.

**Figure S4.**
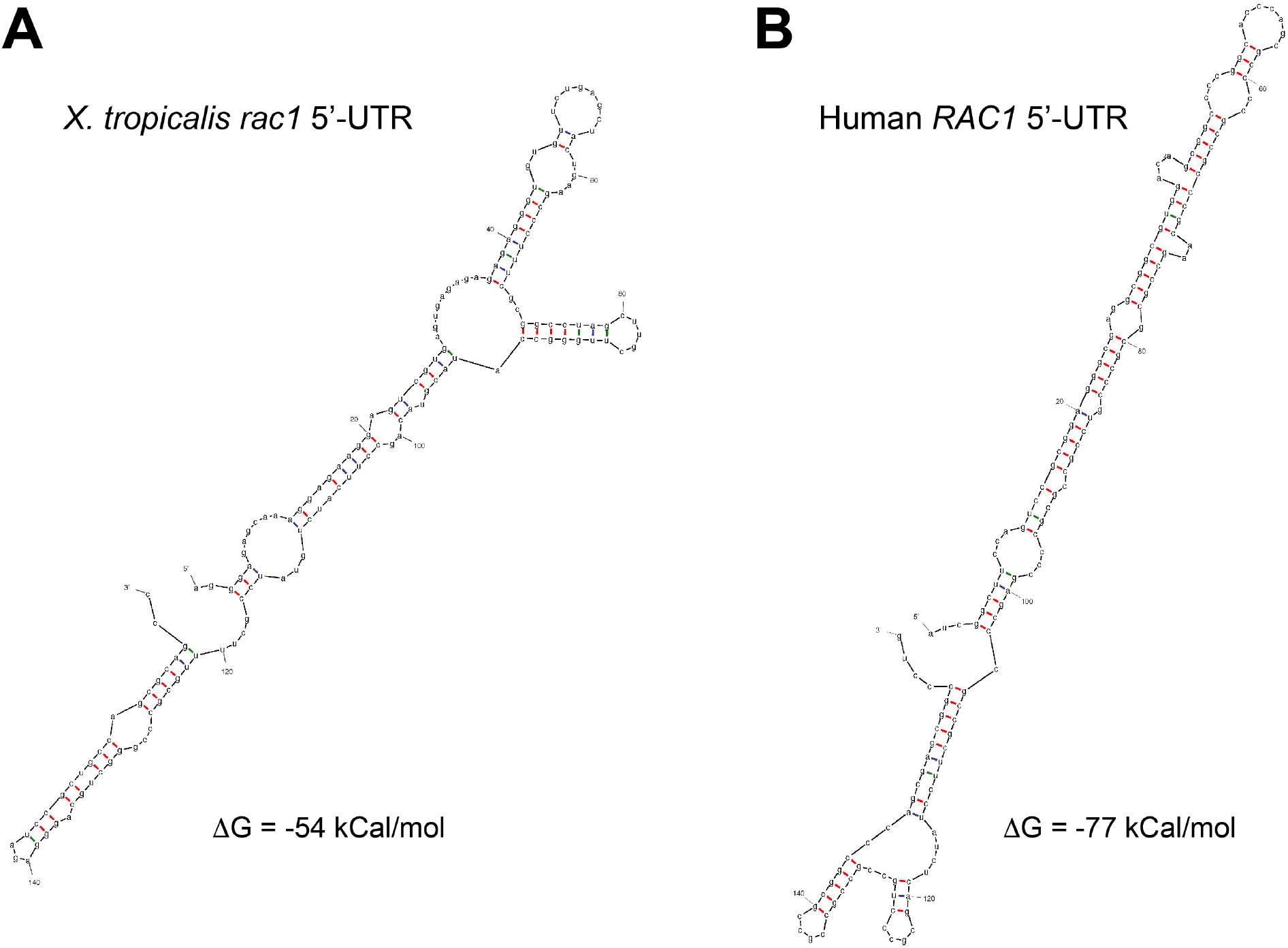
Predicted secondary structures of the *X. tropicalis rac1* and human *RAC1* 5’-UTRs. Prediction was carried out using the Mfold algorithm (http://unafold.rna.albany.edu/?q=mfold/download-mfold).

